# High-throughput phenotyping reveals a link between transpiration efficiency and transpiration restriction under high evaporative demand and new loci controlling water use-related traits in African rice, *Oryza glaberrima* Steud

**DOI:** 10.1101/2021.11.28.470237

**Authors:** Pablo Affortit, Branly Effa-Effa, Mame Sokhatil Ndoye, Daniel Moukouanga, Nathalie Luchaire, Llorenç Cabrera-Bosquet, Maricarmen Perálvarez, Raphaël Pilloni, Claude Welcker, Antony Champion, Pascal Gantet, Abdala Gamby Diedhiou, Baboucarr Manneh, Ricardo Aroca, Vincent Vadez, Laurent Laplaze, Philippe Cubry, Alexandre Grondin

## Abstract

Because water availability is the most important environmental factor limiting crop production, improving water use efficiency, the amount of carbon fixed per water used, is a major target for crop improvement. In rice, the genetic bases of transpiration efficiency, the derivation of water use efficiency at the whole-plant scale, and its putative component trait transpiration restriction under high evaporative demand, remain unknown. These traits were measured in a panel of 147 African rice *Oryza glaberrima* genotypes, known as potential sources of tolerance genes to biotic and abiotic stresses. Our results reveal that higher transpiration efficiency is associated with transpiration restriction in African rice. Detailed measurements in a subset of highly differentiated genotypes confirmed these associations and suggested that the root to shoot ratio played an important role in transpiration restriction. Genome wide association studies identified marker-trait associations for transpiration response to evaporative demand, transpiration efficiency and its residuals, that links to genes involved in water transport and cell wall patterning. Our data suggest that root shoot partitioning is an important component of transpiration restriction that has a positive effect on transpiration efficiency in African rice. Both traits are heritable and define targets for breeding rice with improved water use strategies.

## Introduction

Rice is the staple food for more than half of the world’s population and its consumption is continuously growing. In Africa, rice is mainly cultivated in the Western part of the continent, where its production increased by 104.3 % from 2009 to 2019 (FAOSTAT, 2021). A further increase of 79.4 % will be needed by 2025 to meet the projected local demand (FAOSTAT, 2021). Most of the rice grown worldwide is of *Oryza sativa* L. (Asian rice) type that has high yield potential. In West Africa, improving rice productivity is challenged by the reduction of water resources for agriculture due to dryer and hotter climates and increased competition from cities and industries due to rapid population growth (van Oort and Zwart, 2018). In this context, developing agronomic approaches that reduce water use (e.g. aerobic rice or alternate wet and dry cultivation) and rice varieties with better water use efficiency (WUE) is of major interest.

WUE corresponds to the ratio of plant carbon gain to water use (Leakey *et al*., 2019). Beneath this simple definition are a number of component traits (root water uptake or photosynthesis for instance) and numerous surrogate traits (e.g. carbon isotope discrimination or specific leaf area) making WUE a broad idea that can be conceptualized at multiple scales (Hatfield and Dold, 2019). At the plant scale, WUE is described as transpiration efficiency (TE), i.e. the ratio between biomass (usually shoot biomass) and total water transpired to produce this biomass (Vadez *et al*., 2014). Heritable variations in TE have been observed in a number of species including *Arabidopsis thaliana* (Vasseur *et al*., 2014), sorghum (Vadez *et al*., 2011), groundnut (Vadez and Ratnakumar, 2016) or foxtail millet (Krishnamurthy *et al*., 2016; Feldman *et al*., 2018). In rice, genetic determinants of intrinsic WUE measured through carbon isotope discrimination have been observed (This *et al*., 2010), but although genetic variation in TE exists (Ouyang *et al*., 2017), genetic dissection of TE has not been reported. Enhanced expression of several genes involved in diverse physiological mechanisms such as gibberellin-plant mediated architecture modifications (*OsGA2*; Lo *et al*., 2017), promotion of lateral root initiation (*OsHVA1*; Chen *et al*., 2015), reduced stomatal density (*AtERECTA*; Shen *et al*., 2015) or promotion of photosynthesis assimilation in mesophyll cells (*AtHARDY*; Karaba *et al*., 2007) specifically improved TE under irrigated conditions in rice, highlighting the complexity of this trait.

Transpiration restriction is another physiological mechanism that can improve TE (Sinclair *et al*., 2017). For the plant, this strategy translates into opening its stomata and maximizing C assimilation when the vapor pressure deficit (VPD) in the air is below a certain threshold (usually between 1.5 to 2.5 kPa), and closing its stomata when VPD exceeds this threshold resulting in lower stomatal conductance and consequently reducing water use (Condon, 2020). Transpiration restriction can be measured by the slope of the transpiration response to increasing VPD or by the inflexion point in transpiration response (usually inversely correlated with the slope). Large genetic variations in transpiration response to increasing VPD have been observed in maize (Gholipoor *et al*., 2013), wheat (Medina *et al*., 2019), sorghum (Choudhary and Sinclair, 2014; Choudhary *et al*., 2020) or pearl millet (Kholová *et al*., 2010), suggesting that this response is determined by genetic factors (Vadez *et al*., 2014; Sinclair *et al*., 2017). In pearl millet, transpiration restriction was associated with terminal drought tolerance and quantitative trait loci (QTLs) controlling both traits were found to colocalize (Kholová *et al*., 2010, 2012). Although transpiration restriction can lead to reduction in biomass as observed in wheat plants grown under irrigated environments (Medina *et al*., 2019), it can also lead to soil water conservation at vegetative stage and improve yield in pearl millet plants grown under drought stress (Vadez *et al*., 2013). Transpiration restriction could therefore be an interesting trait to deploy for improving TE and drought tolerance of upland rice grown under dry, hot and drought-prone environments of West Africa.

African rice *O. glaberrima* Steud. was domesticated in the inner Niger delta from a wild Sahelian ancestor *O. barthii* A. Chev. (Cubry *et al*., 2018). It is adapted to dry environments and has raised the interest of the scientific community because of its potential reservoir of tolerance genes to biotic and abiotic stresses (Wang *et al*., 2014). Recently, high-depth re-sequencing data of 163 *O. glaberrima* genotypes originating from diverse environments in West Africa was used to infer the origin of domestication of African rice (Cubry *et al*., 2018). This panel was further used to identify QTLs associated with flowering time, panicle architecture and resistance to *Rice yellow mottle virus*, providing insights into the adaptive variation of African rice as compared to Asian rice (Cubry *et al*., 2020). Due to its adaptation to contrasted environments, we hypothesized that *O. glaberrima* could also be a source of interesting alleles for improving TE. Here, we phenotyped 147 fully sequenced *O. glaberrima* genotypes for shoot growth and water use dynamics to derive transpiration restriction, TE and its residuals at 29 days after sowing. A subset of contrasted genotypes for TE were further studied to better understand the physiological determinants of these complex traits and genome wide association studies (GWAS) allowed the dissection of their genetic bases.

## Material and methods

### Plant material

A panel of 147 fully sequenced traditional cultivated genotypes of African rice (*O. glaberrima*) that were sampled from 1974 to 2005 mainly in West Africa, with few genotypes from East Africa was used in this study (Cubry *et al*., 2018).

### Plant growth conditions and measurements

#### Large-scale phenotyping experiment

Large-scale phenotyping of shoot growth and water use in the *O. glaberrima* panel was performed using the PhenoArch platform hosted at M3P, Montpellier Plant Phenotyping Platforms (https://www6.montpellier.inra.fr/lepse/M3P) located at INRAE Montpellier (43°37’03.6”N; 3°51’27.9”E). Dehusked seeds were sown in biodegradable tray pots (55% white peat and 45% woodpulp, pH 5.0; Jiffy) containing a mix of fine clay (20%) and fine, blond and black (30, 10 and 40%, respectively) peats at pH 6, and amended with 1.5kg of 14-16-18 N-P-K for 25 L (Substrat SP 15%, KLASMANN). From sowing to 15 days after sowing (DAS), plants were grown under irrigated conditions in a greenhouse at the Institut de Recherche pour le Développement (IRD) in Montpellier (43°38’41.31”N; 3°51’57.3”E) under 12h photoperiod, day temperature of 28°C, night temperature of 25°C and with 60-70% humidity. Fifteen days after sowing, plants were transferred to the PhenoArch greenhouse and individual seedlings in their biodegradable pot were transplanted into 9L pots filled with the same soil. Plants were exposed to the same environmental conditions as in IRD facilities and grown for two more weeks under irrigated conditions (soil water potential maintained at −0.05 MPa). The experiment was arranged in a randomized complete block design with seven replications.

The PhenoArch platform is structured as a conveyor belt feeding the plants to imaging or watering units as described in Cabrera-Bosquet *et al*., (2016). The shoot imaging unit is composed of three cabins equipped with top and side RGB cameras and LED illumination. The watering units are composed of five weighing terminals and high-precision pumps that allow monitoring of the soil water content. Imaging and watering routines were sequentially performed every day from 18 to 29 DAS. Plants were further moved back to the same positions and orientation in order to keep position throughout the experiment.

Shoot biomass and leaf area were estimated every day from images. Briefly, RGB images were taken for each plant from 13 views (12 side views from 30° rotational difference and one top view) and pixels from each image were extracted from those of the background as described in Brichet *et al*. (2017). Whole plant leaf area and shoot fresh weight were then estimated using calibration curves built using multiple linear regression models based on processed images against ground truth measurements of leaf area and fresh biomass (Supplementary Fig. S1).

Leaf area and daily water lost by the pot was used to measure transpiration rate. Plant daily water uptake from 18 to 29 DAS was added to calculate total water uptake. Transpiration efficiency (TE) was measured as the ratio between shoot fresh weight and total water uptake at 29 DAS. Because shoot mass is intrinsically related to plant water use, the residuals of TE (TEr) were calculated as the genotype-specific deviation from the least squares linear regression between total water uptake and shoot fresh weight measured at 29 DAS. Reference evapotranspiration was calculated according to the Penman-Monteith formula (Zotarelli *et al*., 2014), as a proxy for the evaporative demand. Averaged transpiration rate was then plotted against maximum reference evapotranspiration for five windows of time, i.e. at 23, 25, 26, 27 and 28 DAS for each genotype and the slope of the corresponding linear regression was calculated to evaluate transpiration response to evaporative demand (SlopeTR).

### Small-scale phenotyping experiment

A sub-set of ten genotypes contrasting for shoot biomass (Og_118, Og_12, Og_162, Og_61, Og_62, Og_15, Og_184, Og_185, Og_408 and Og_43) was grown at IRD in 5.5L pots containing a potting mixture (M2 substrate, Jiffy) receiving optimal fertilization and under the same environmental conditions as above with five replications per genotypes. Pots were randomized and soil was well irrigated and covered with 2-3 cm of plastic beads to prevent soil evaporation. At 29 DAS, pots were transferred into an adjacent greenhouse on top of balances monitoring weight every 30 min (Phenospex Ltd). Vapor pressure deficit (VPD) was monitored and reached values around 4-5 KPa around noon. At 36 DAS, leaves were harvested to measure leaf area using a planimeter (LI-3100C, LI-COR), shoot fresh weight and tiller number. Plant water uptake from 29 to 36 DAS was used to calculate total water uptake. The root system was carefully washed from the soil and placed, along with fresh shoots, into an oven for 3 days at 60°C to measure root dry weight, shoot dry weight and root to shoot ratio. TE and TEr were calculated from shoot fresh weight and total water uptake at 36 DAS as in the large-scale experiment. Transpiration profiles and features characterizing these profiles were obtained from an adapted automated pipeline developed by (Kar *et al*., 2020). Leaf area measured at 36 DAS was further used to measure transpiration rate at 35 DAS, assuming marginal changes in leaf area between 35 and 36 DAS. Averaged transpiration rate for each genotypes were plotted against time between 9 AM to 4 PM, as a proxy for the evaporative demand, and the segmented R package v1.2.0 (Muggeo, 2008) was used to calculate the slope of the initial linear regression and an inflexion point in the transpiration response.

### Data analysis

A non-parametric smoothing approach was used to detect outliers in the time-course shoot fresh weight and water uptake from the large-scale phenotyping experiment (Millet *et al*., 2021). This approach uses a *locfit()* function that fits a local regression at a set of points and a *predict()* function to interpolate this fit to other points. A confidence interval is calculated and points located outside the interval are considered as outliers.

Shoot fresh weight and total water uptake datasets were further analyzed for detection of plant outliers. For this, a multi-criteria analysis with expert rules function was used (Millet *et al*., 2021). Leaf area, shoot fresh weight and plant height were modelled considering fixed experiment effects, and random genotypic, replicate and spatial effects using SpATS. A lower and an upper bound interval for the evolution of these traits across the experiment was created and plants with lesser shoot fresh weight than the lower bound interval were considered as small outlier plants while plants with greater shoot fresh weight than the upper bound interval were considered as large outlier plants. These plants were removed from the dataset.

Shoot fresh weight and total water uptake values were finally corrected for spatial heterogeneity in the PhenoArch greenhouse using the StatgenHTP R package (Millet *et al*., 2021). This package is based on the previously developed SpATS (Spatial Analysis of Trials using Splines; Rodríguez-Álvarez *et al*., 2015) package and separate the genetic effect from the spatial effect by taking into account the evolution of shoot fresh weight and total water uptake across time.

### Genome wide association studies

The *O. glaberrima* panel used in this study was previously subjected to in-depth re-sequencing to identify Single Nucleotide Polymorphisms (SNPs) based on mapping to the *O. sativa japonica* cv. Nipponbare reference genome MSU7 (Cubry *et al*., 2018). A total of 892,539 SNPs was identified for this panel with a genome-wide high linkage disequilibrium at short distance (0.2 for at least 150 kb) that slowly decayed (Cubry *et al*., 2020). Missing data remaining in the SNP matrix (less than 5% of missing data per SNP) were imputed using the *impute* function of the LEA R package v3.1.0 (Frichot and François, 2015; Gain and François, 2021).

Association between genomic polymorphisms and mean phenotypic variables were performed using a pipeline described in Cubry *et al*. (2020). In this pipeline, SNPs displaying a minimal allele frequency (frequency of the minor allele) lower than 5% are filtered out. A simple non-corrected linear model (analysis of variance, ANOVA) was performed to assess the effect of confounding factors such as relatedness and population structure on false positive rates. Two genome-wide association methods were further used: (1) a latent factor mixed model as implemented in the LFMM v2 R package that jointly estimated associations between genotype and phenotype and confounding factors (Frichot *et al*., 2013); and (2) an efficient mixed model analysis (EMMA) as implemented in the EMMA R package (Kang *et al*., 2008). Population structure was corrected using four latent factors in the LFMM model and a similarity-based kinship matrix in the EMMA model (Cubry *et al*., 2020). The results of all analyses were graphically represented as Manhattan plots and QQ-plot to assess efficiency of confounding factors correction using the qqman R package v0.1.4 (Turner, 2018). A *p*-value threshold of 10^−5^ was used to select significant SNPs. Candidate genes were selected in a 150kb region upstream and downstream of the significant SNPs by intersecting the region with the MSU7 genome annotation (Kawahara *et al*., 2013).

### Statistical analyses

Statistical analyses were performed with R version 4.0.2 (R Development Core Team, 2018) using ANOVA (aov function) to detect genotypic and environmental effects. To determine to which extent the measured traits were genetically determined, broad-sense heritability (*H*^2^) was calculated according to (Oakey *et al*., 2006) using the *inti* R package v0.4.3 (Lozano-Isla, 2021) according to the following formula:

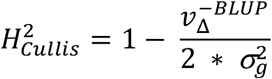

where σ^2^ refers to the variance, *g* to the genotype, 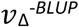 to the average standard error of the genotypic best linear unbiased prediction (BLUP).

## Results

### Water-use related traits are highly variable and heritable in African rice

In order to measure variation in water-use related traits in *O. glaberrima*, 147 genotypes from a diversity panel were grown in the INRAE-PhenoArch high-throughput phenotyping platform where shoot growth and water uptake were monitored daily by imaging and pots weighing from 17 to 29 DAS. These data were used to calculate plant leaf area, water uptake, transpiration rate and shoot fresh weight during the course of the experiment. Moreover, shoot fresh weight was measured at the end of the experiment (29 DAS). Shoot biomass accumulation over time was accompanied by an increase in water uptake and better discrimination between genotypes (Fig. 1A and B). Large genotypic variation in shoot fresh weight and total water uptake was observed at 29 DAS (coefficient of variation of 31.68 % and 10.17 %, respectively; Table 1). Transpiration efficiency (TE) calculated from shoot fresh weight and total water uptake at 29 DAS ranged from 1.82 to 11.35 and showed a coefficient of variation of 23.89 % (Table 1 and Fig. 1C). Residuals representing the genotype-specific deviation from the least squares linear regression between total water uptake and shoot fresh weight measured at 29 DAS were calculated and named TEr (Supplementary Fig. S2). TEr represents the genotype-specific component of shoot biomass that is independent of water uptake or, in other words, the difference of shoot biomass produced for the same amount of water consumed. TEr varied from −5.24 to 3.34 in the panel (Table 1). Except for transpiration rate, all traits were subjected to significant genotypic effect resulting in high heritability (> 0.9 for shoot fresh weight, total water uptake and TE and 0.7 for Ter; Supplementary Tables S1, S2, S3, S4, S5 and Supplementary Fig. S3, S4, S5, S6, S7).

**Table 1:** Variation of water-use related traits in the *O. glaberrima* panel. Range, mean, standard deviation (SD), coefficient of variation (CV) and broad-sense heritability (*H*^2^) for shoot fresh weight, total water uptake, transpiration efficiency and residuals of transpiration efficiency were measured at 29 days after sowing. Transpiration rate was plotted against maximum reference evapotranspiration at 23, 25, 26, 27 and 28 days after sowing and the slope of the corresponding linear regression was used to estimate transpiration response to evaporative demand (SlopeTR).

| Trait | Description | Range | Mean | SD | CV | $H^2$ |
| --- | --- | --- | --- | --- | --- | --- |
| SFW | Shoot fresh weight (g) | 1.28-14.24 | 6.71 | 2.12 | 31.68 | 0.92 |
| TWU | Total water uptake (g) | 704.26-1382.86 | 999.1 | 101.6 | 10.17 | 0.88 |
| TE | Transpiration efficiency (g SFW g<sup>-1</sup> TWU) | 1.82-11.35 | 6.60 | 1.58 | 23.89 | 0.91 |
| TEr | Residuals of transpiration efficiency | -5.24-3.34 | 0.0009 | 1.09 | 126167 | 0.73 |
| TR | Transpiration rate (ml water sec<sup>-1</sup> cm<sup>-2</sup>) | 0.01-0.05 | 0.03 | 0.006 | 18.71 | 0.16 |
| SlopeTR | Transpiration response to increasing evaporative demand | 0.2-3.26 | 0.69 | 0.24 | 34.71 | 0.42 |

**Fig. 1:**
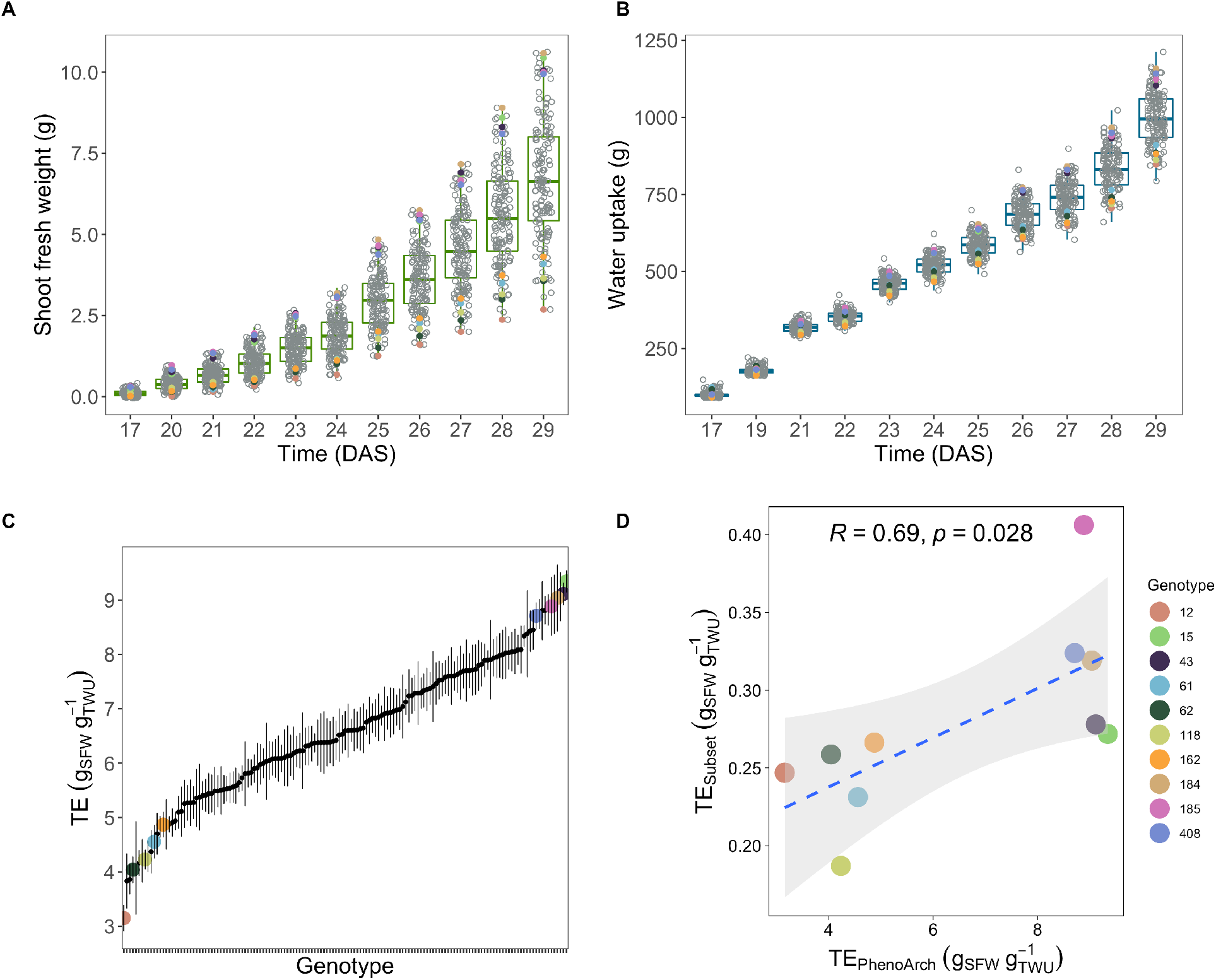
Variation in shoot fresh weight, water uptake and transpiration efficiency (TE) in *O. glaberrima*. A-B: Variation in shoot fresh weight (A) and water uptake (B) from 17 to 29 days after sowing (DAS) during the large-scale experiment. C: Variation in TE measured as the ratio between shoot fresh weight and total water uptake at 29 DAS in the large-scale experiment. D: Covariation between TE measured in the large-scale experiment (PhenoArch) and in a subset of genotypes in the small-scale experiment. R: Pearson’s correlation coefficient; *p*: *p*-value of the Pearson’s correlation test; SFW: shoot fresh weight; TWU: total water uptake. Genotypes from the subset are highlighted in A, B and C following the same color legend as in D.

In order to check the repeatability of the traits measured in the large-scale phenotyping experiment, a subset of ten genotypes contrasting for shoot fresh weight, total water uptake and TE were selected and grown in the small-scale experiment for measurements of the same variables at 36 DAS. Genotypes Og_118, Og_12, Og_162, Og_61 and Og_62 had low shoot fresh weight, total water uptake and TE while genotypes Og_15, Og_184, Og_185, Og_408 and Og_43 had high shoot fresh weight and total water uptake and TE (Supplementary Fig. S8). Shoot fresh weight and total water uptake were significantly positively correlated (*p*-value < 0.01) between the two experiments showing their robustness between environments (Supplementary Fig. S9). Similarly, significant correlations between TE measured in the large-scale and small-scale experiments were observed (Fig. 1D).

Since transpiration response to increasing evaporative demand contributes to modulate TE, we calculated the slope of the regression between transpiration rate and maximum reference evapotranspiration at 23, 25, 26, 27 and 28 DAS in the large-scale experiment (Supplementary Fig. S10). The slope of the linear regression (SlopeTR) represents the transpiration response to increasing evaporative demand and reads as follows: the lower the slope, the lower the genotype responds to the evaporative demand by restricting its transpiration. SlopeTR showed a coefficient of variation of 34.71 % in the population and a broad-sense heritability of 0.42 (Table 1 and Supplementary Fig. S11).

Overall, our data show that water use-related traits were variable and highly heritable in the *O. glaberrima* panel. These variations appeared to be conserved in a subset of genotypes across environments.

### Transpiration restriction under high evaporative demand contributes to increased TE in African rice

To better understand the relationship between the different water use-related traits, we performed correlation analyses between each trait measured in the *O. glaberrima* panel. A significant positive correlation was observed between shoot fresh weight (SFW) and total water uptake (TWU; *p*-value < 0.001; Fig. 2A) indicating that plants that grew a bigger biomass used more water. Similarly, a positive significant correlation was observed between TE and both shoot fresh weight and total water uptake (*p*-value < 0.001). On the other hand, TEr was correlated with TE (R = 0.73) and less so with shoot fresh weight and total water uptake, although all correlations were significant (*p*-value < 0.001; Fig. 2A). Interestingly, a significant negative correlation was observed between TE and transpiration response to evaporative demand (SlopeTR; Fig. 2B), suggesting that genotypes with lower transpiration at higher evaporative demand had higher TE. This indicates that transpiration restriction under high evaporative demand might contribute to increased TE in African rice.

**Fig. 2:**
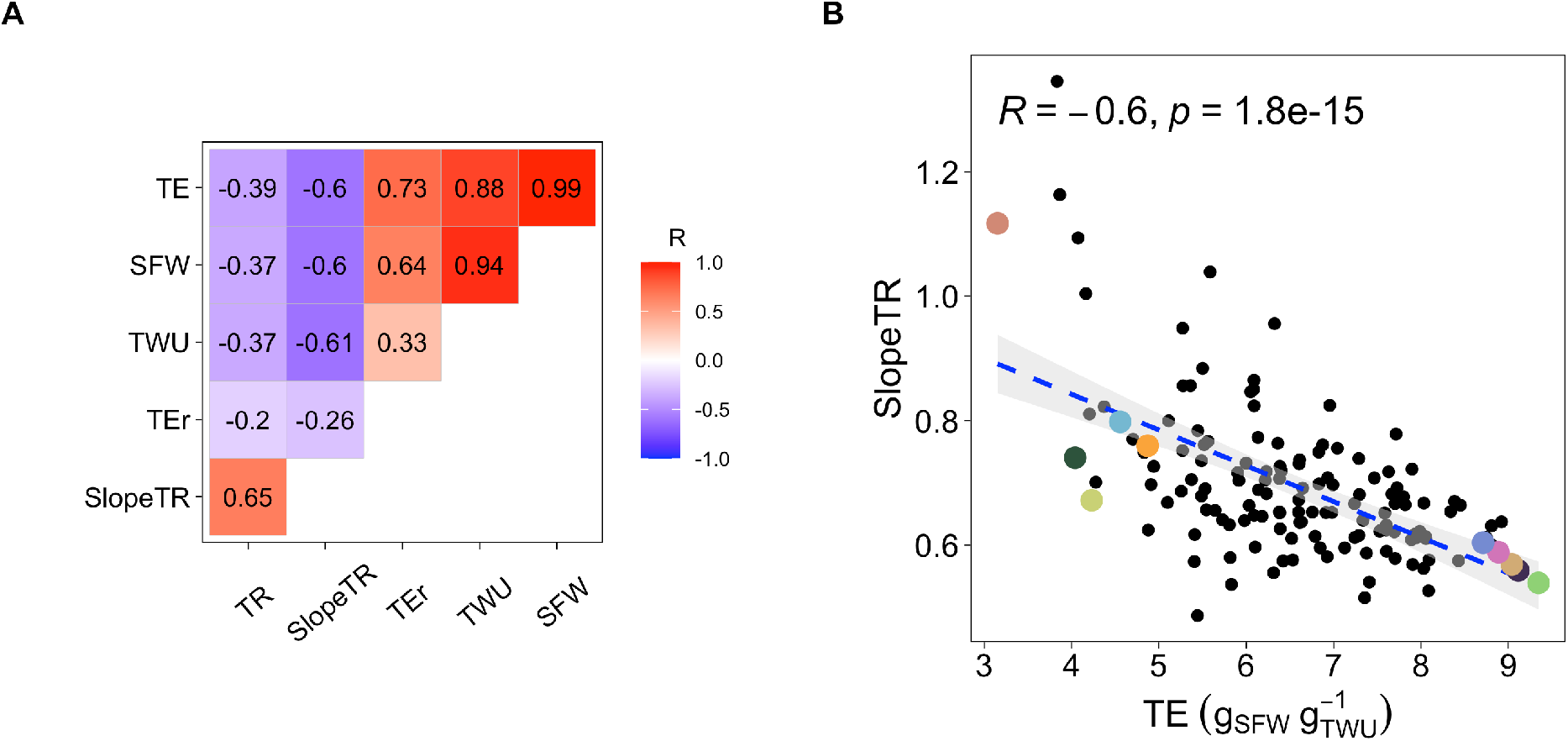
Correlation between water use-related traits in *O. glaberrima*. A: Pearson’s correlation coefficient (R) between averaged values shoot fresh weight (SFW), total water uptake (TWU), transpiration efficiency (TE), residuals of transpiration efficiency (TEr) at 29 days after sowing and transpiration response to evaporative demand (SlopeTR) measured in the large-scale experiment. B: Covariation between TE and SlopeTR. Dots represent the averaged TE plotted against the average SlopeTR for individual genotypes. Genotypes highlighted in color are from the subset used in the small-scale experiment, following the same color legend as in Fig. 1.

To test this hypothesis, we studied the transpiration response to VPD across one day in the same subset of contrasted genotypes as above at 35 DAS. During this experiment, VPD gradually increased to reach its maximum around 5 PM and further decreased (Fig. 3A). Transpiration rate followed the same pattern until 4 PM where it reached its maximum for most of the genotypes and further decreased (Fig. 3A). Large variation in the transpiration response to VPD was observed among genotypes, with Og_185 and Og_118 having the lowest and highest maximum transpiration rate at 4 PM (Fig. 3A). To further quantify this response, we measured the slope of transpiration response to increasing VPD and the time of transpiration inflexion between 10 AM to 4:30 PM (Supplementary Fig. S12). While the inflexion time did not significantly vary among genotypes, significant differences were observed in the slope of transpiration response (SlopeTR; *p*-value < 0.01; Supplementary Table S7). Principal component analyses showed that the first principal component that separated the genotypes with low shoot fresh weight from those with high shoot fresh weight accounted for 73.3 % of the global variation (Fig. 3B). Genotypes with low shoot fresh weight covaried with transpiration rate and transpiration response to evaporative demand (SlopeTR) while genotypes with higher shoot fresh weight covaried with root dry weight, total water uptake and TE (Fig. 3B). Transpiration response to evaporative demand (SlopeTR) was significantly negatively correlated with shoot fresh weight (*p*-value < 0.05), total water uptake (*p*-value < 0.01) and TE (*p*-value < 0.05; Fig. 3C and Supplementary Fig. S13). Interestingly, the ratio of root to shoot dry weight (Root:Shoot ratio) was significantly positively correlated with transpiration response to evaporative demand (SlopeTR; *p*-value < 0.05; Fig. 3D) and tended to covary with transpiration rate (R = 0.4), although this covariation was not significant (Supplementary Fig. S13).

**Fig. 3:**
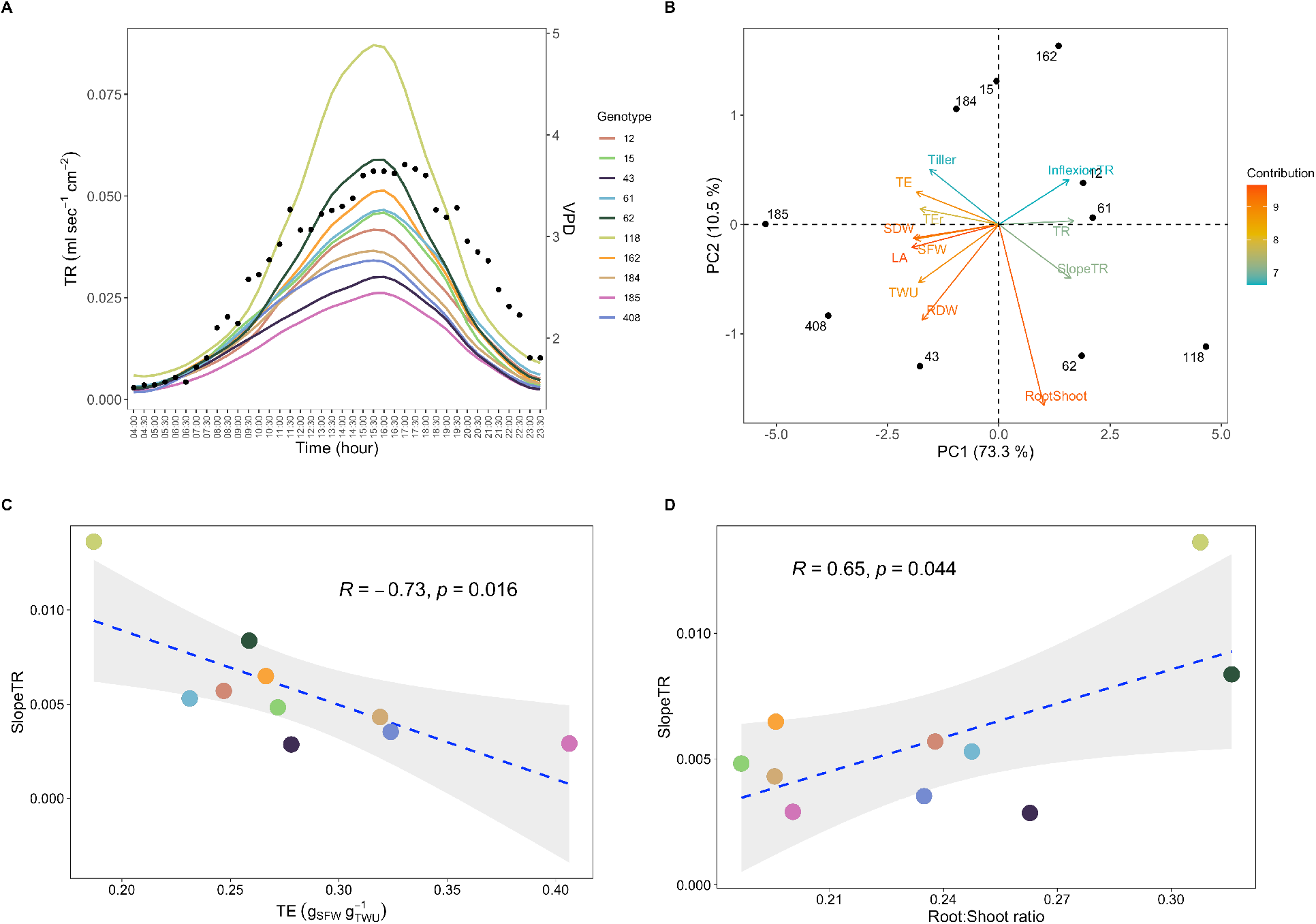
Relation between water use-related traits and plant morphology in a subset of *O. glaberrima* genotypes. A: Evolution of transpiration rate (TR; colored lines) upon increasing vapor pressure deficit (VPD; black dots) measured in the small-scale experiment at 35 days after sowing. B: Relationship between tiller number (Tiller), transpiration efficiency (TE), residuals of transpiration efficiency (TEr), shoot fresh weight (SFW), shoot dry weight (SDW), leaf area (LA), total water uptake (TWU), root dry weight (RDW), root to shoot ratio (RootShoot) transpiration response to increasing evaporative demand (SlopeTR), TR and TR inflexion in response to increasing evaporative demand (InflexionTR) measured at 35 days after sowing in the small-scale experiment, and analyzed using principal component (PC) analysis. C-D: Covariation between TE and SlopeTR (C) and between root to shoot ratio and SlopeTR (D). Dots represent the averaged TE or root to shoot ratio plotted against the average SlopeTR for individual genotypes. R: Pearson’s correlation coefficient; *p*: *p*-value of the Pearson’s correlation test. Color legend is the same as in A.

Altogether, precise measurements of transpiration under increasing evaporative demand in the subset of genotypes confirmed that transpiration restriction to increasing evaporative demand (lower SlopeTR) was associated with higher TE in African rice. Our results further demonstrated that shoot biomass and the balance between roots and shoots growth played important roles in the transpiration response to increasing evaporative demand.

### Identification of genomic regions associated with water use-related traits by association genetics

As our data showed that water use-related traits were variable and highly heritable in the *O. glaberrima* panel, we performed association genetics to identify polymorphisms associated with their variation. Genomic regions associated with shoot fresh weight, total water uptake, TE, its residuals TEr and transpiration response to increasing evaporative demand (SlopeTR) were identified using two GWAS methods (LFMM and EMMA). Applying a 10^−5^ *p*-value threshold, we observed a total of 42, 59, 49, 95 and 29 significant marker-trait associations in both methods for shoot fresh weight, total water uptake, TE, TEr and transpiration response to evaporative demand (SlopeTR), respectively (Fig. 4A–D and 5A, Supplementary Fig. S14A–D and S14A). The two methods allowed an efficient correction for false positives linked to genetic structure (QQ-plots, Fig. 4E–F and 5B, Supplementary Fig. S14E–F and S15B).

**Fig. 4:**
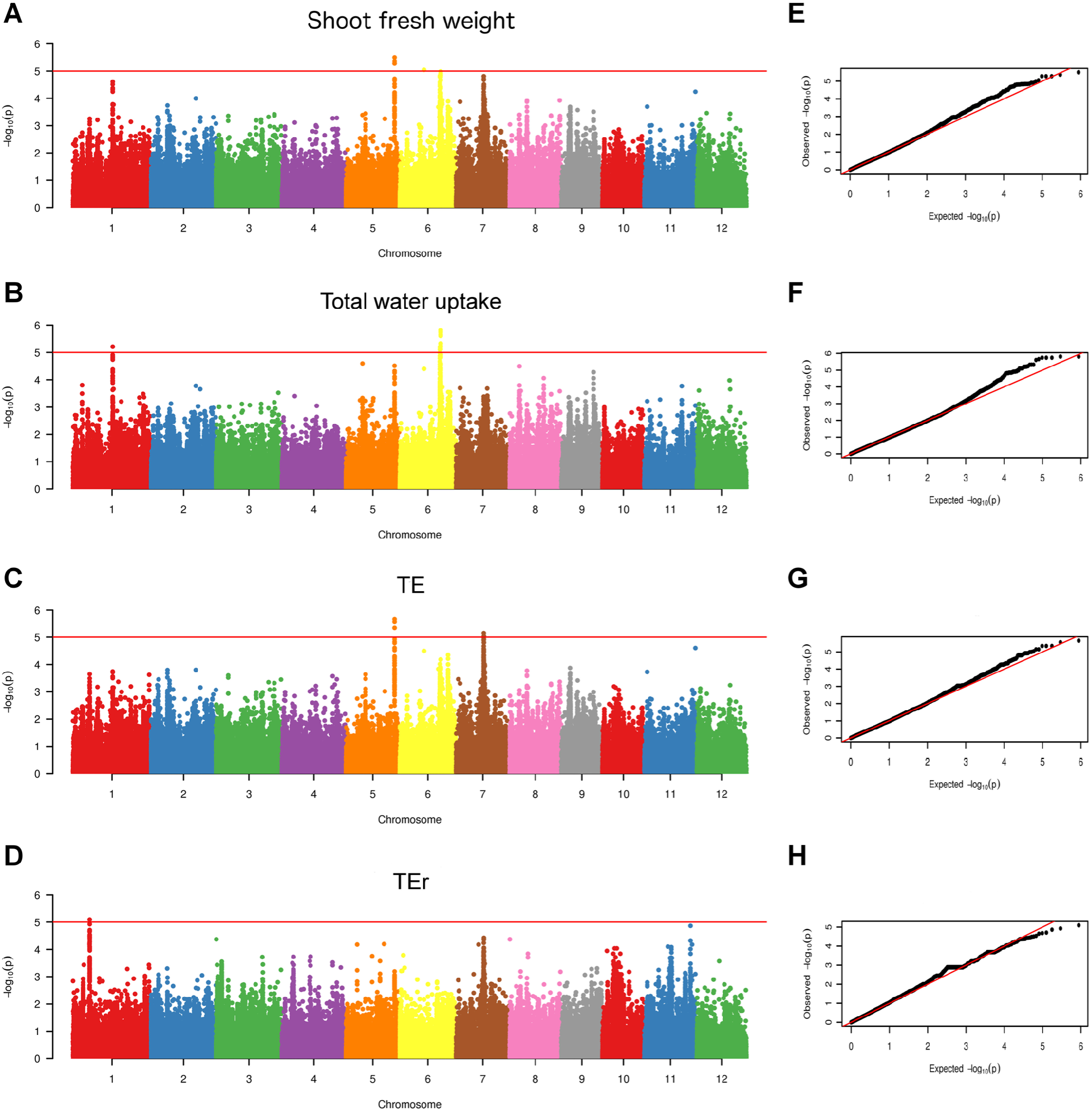
Genome wide association studies for shoot fresh weight, total water uptake, transpiration efficiency (TE), and residuals of transpiration efficiency (TEr) in *O. glaberrima*. Manhattan plots (A-D) and QQ-plots (E-H) obtained with the latent factor mixed model (LFMM) are shown. Manhattan plots show the log10 *p*-value (y axis) at each SNP position on the different chromosomes (x axis). The red lines in A-D delimit the threshold for highly significant SNPs (*p*-value < 10^−5^).

**Fig. 5:**
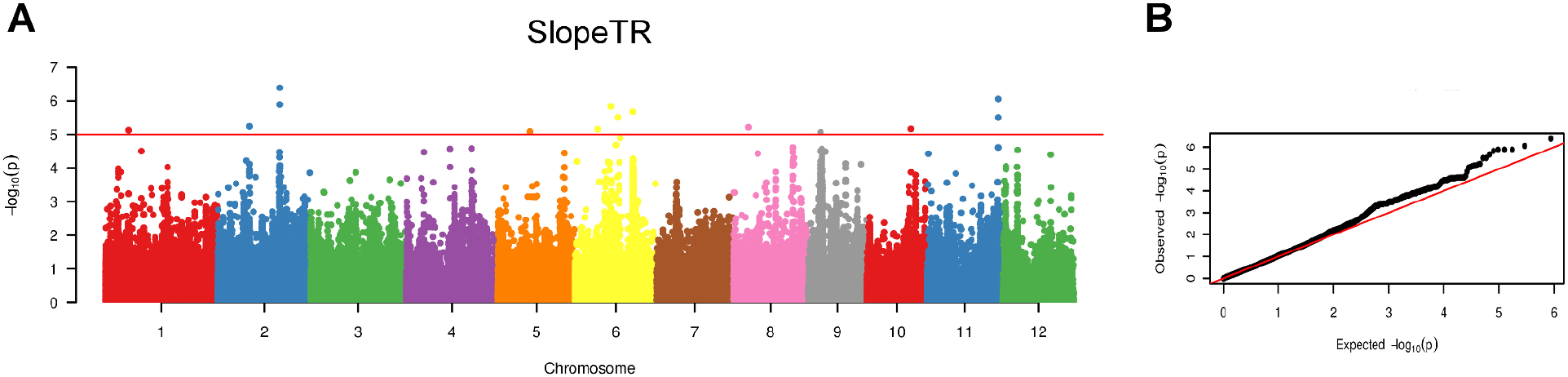
Genome wide association studies for transpiration response (SlopeTR) to increasing evaporative demand in *O. glaberrima*. Manhattan plots (A) and QQ-plots (B) obtained with the latent factor mixed model (LFMM) are shown. Manhattan plots show the log10 *p*-value (y axis) at each SNP position on the different chromosomes (x axis). The red line in A delimits the threshold for highly significant SNPs (*p*-value < 10^−5^).

Significant associations were observed for shoot fresh weight and TE on chromosome 5, and for shoot fresh weight and total water uptake on chromosome 6 using both LFMM and EMMA methods (Fig. 4 and Supplementary Fig. S14). Further significant associations were observed with LFMM on chromosome 1 for total water uptake (-log_10_ (*p*-value) = 4.63 in EMMA; Fig. 4B and Supplementary Fig. S14B) and on chromosome 7 for TE (-log_10_ (*p*-value) = 3.89 in EMMA; Fig. 4C and Supplementary Fig. S14C). Although not significant, similar associations were observed on chromosome 1 for shoot fresh weight and chromosome 7 for shoot fresh weight and TEr in LFMM (Fig. 4A and D). More specific associations were observed for TEr on chromosome 1 in LFMM and EMMA and on chromosome 4 in EMMA (-log_10_ (*p*-value) = 3.73 in LFMM; Fig. 4D and Supplementary Fig. S14D). For transpiration response to evaporative demand (SlopeTR), the two most significant associations were observed on chromosome 2 and 11 in both LFMM and EMMA (Fig. 5A and Supplementary Fig. S15A).

Interestingly, the GWAS association located on chromosome 5 position 26971730 for shoot fresh weight and TE co-localized with a previously reported QTL for early vigor in Asian rice (*Oryza sativa*; Cui *et al*., 2002). Two alleles were present for the corresponding SNP with plants carrying either an adenine (A; 45.5 % of the panel) or a guanidine (G; 54.5 % of the panel), with plants carrying the G allele having a 25.2 % shoot biomass gain (Supplementary Fig. S16). Genotypes with low shoot fresh weight (Og_12, Og_61, Og_62, Og_118 and Og_162) grown in the small-scale experiment carried the A allele while genotypes with high shoot fresh weight (Og_15, Og_43, Og_184, Og_185, Og_408) carried the G allele (Supplementary Fig. S8), confirming the allelic distribution observed in the large-scale experiment. Hence, our results confirmed the importance of this genomic region to control early shoot growth and the conservation of this QTL in Asian and African rice.

Altogether, our association genetics approach identified at least 14 potential genetic regions associated with water use related-traits in African rice, some of which are specific to TE, TEr or transpiration response to evaporative demand.

### Genes potentially involved in photosynthesis, regulation of water transport and drought responses are underlying associations for water use-related traits

We next analyzed the genes present in genetic regions associated with shoot fresh weight, total water uptake, TE, TEr and transpiration response to evaporative demand. Since linkage disequilibrium is high at short distance then slowly decays to values below 0.2 after around 150 kb in this panel (Cubry *et al*., 2018, 2020), we considered genes in a 300 kb region surrounding the most significant SNP at each association peak (Table 2). The most significant SNP on chromosome 5 position 26971730 associated with shoot fresh weight and TE mapped in an intergenic region at 1.8 Kb of a gene encoding a Polyprenyl Synthetase and 10.2 Kb of a gene encoding a Ras-related nuclear protein (RAN) GTPase-activating protein. Polyprenyl synthetase catalyzes the synthesis of isopentenyl diphosphate that is involved in the biosynthesis pathway of plastoquinones, essential proteins for the photosynthesis machinery carrying electrons in the linear and alternative electron chains (Liu *et al*., 2019; Havaux, 2020). RAN GTPase are involved in nucleocytoplasmic transport, mitotic progression and membrane trafficking, cytoskeletal organization or cell polarity, and have important roles in plant growth, development and response to stress conditions (Nielsen, 2020). In particular, the Arabidopsis *RanBP1c* and wheat *Ran1* are involved in auxin-induced root growth and development through the control of mitotic progress (Kim *et al*., 2001; Wang *et al*., 2006). Heterologous overexpression of a wheat Ran GTPase in rice reduced the number of lateral roots and induced hypersensitivity to auxin (Wang *et al*., 2006).

**Table 2:** List of candidate genes identified in genetic regions associated with shoot fresh weight (SWF), total water uptake (TWU), transpiration efficiency (TE), residuals of transpiration efficiency (TEr), and transpiration response to evaporative demand (SlopeTR).

| Trait | Chr_position | Locus (MSU) | Annotation | Hypothetical function | Reference |
| --- | --- | --- | --- | --- | --- |
| SFW, TE | Chr5_26971730 | LOC_Os05g46560 | RAN GTPase-activating protein 1 | Auxin-mediated root development | Kim *et al.*, 2001<br>Wang *et al.*, 2006 |
|  |  | LOC_Os05g46580 | Polyprenyl synthetase | Photosynthesis through Plastoquinone biosynthesis | Liu *et al.*, 2019<br>Havaux, 2020 |
| TWU | Chr1_21989359 | LOC_Os01g39110 | ZOS1-10 C2H2 zinc finger family protein | Development, drought response | Agarwal *et al.*, 2007<br>Huang *et al.*, 2009 |
| TE | Chr7_15311728 | LOC_Os07g26630<br>LOC_Os07g26660<br>LOC_Os07g26690 | Plasma membrane aquaporins | Water transport and use, response to abiotic stresses | Sakurai *et al.*, 2005<br>Guo *et al.*, 2006 |
|  |  | LOC_Os07g26720 | OsRR7, type-A response regulator | Abiotic responses through abscisic acid and cytokinin signaling | Huang *et al.*, 2018<br>Zeng *et al.*, 2021 |
| TEr, SlopeTR | Chr1_9237408 | LOC_Os01g16220 | Sad1 - UNC-like C-terminal domain | Plant development | Li *et al.*, 2015 |
| | | LOC_Os01g15979 | Gβ protein | Cell expansion and lipid metabolism | Choudhury *et al.*, 2019 |
| TEr | Chr4_6544212 | LOC_Os04g11830 | TCP family transcription factor | Leaf development | Koyama *et al.*, 2010<br>Li *et al.*, 2020 |
|  |  | LOC_Os04g12010 | Glycosyltransferase | Cell wall formation | Anders *et al.*, 2012<br>Lin *et al.*, 2016 |
| SlopeTR | Chr2_23902475 | LOC_Os02g39590 | GDSL-like lipase/acylhydrolase | Growth and development, stress responses | Zhang *et al.*, 2017 |

The GWAS peak for total water uptake on chromosome 1 (position 21989359) is located in an intergenic region at 3.5 kb of a gene coding the C-type C2H2 zinc finger protein ZOS1-10 (Table 2). C2H2-type zinc finger proteins form a large family of 189 members in rice (Agarwal *et al*., 2007) having many roles in plant growth, development, abiotic and biotic resistances (Han *et al*., 2020). The rice C2H2 zinc finger protein *Drought and Salt Tolerance* (*DST*) is for instance involved in leaf morphology and water use through stomatal control, its loss of function increasing rice and salt tolerance (Huang *et al*., 2009).

The most significant SNP on chromosome 7 associated with TE mapped on the 3’ UTR of a gene coding an unknown protein (LOC_Os07g26595). A cluster of four Plasma membrane Intrinsic Proteins (PIPs) primarily expressed in roots are located from 47 to 98 Kb away from this SNP (Sakurai *et al*., 2005; Guo *et al*., 2006). Plasma membrane aquaporins are water channels located on the plasma membrane and were described as important contributors of root radial water transport (Grondin *et al*., 2016) and water use efficiency in rice (Nada and Abogadallah, 2014). Another interesting gene coding for OsRR7, a type-A response regulator is located at 130 Kb from this SNP. In Arabidopsis, a type-A response regulator protein ARR5 is phosphorylated by SnRK2s protein and amplifies the ABA-mediated stress response while inhibiting the cytokinin-responsive genes promoting growth and development (Huang *et al*., 2018). In maize, a recent report demonstrated that natural variation in type-A response regulator confers chilling tolerance (Zeng *et al*., 2021).

The GWAS associations for TEr and transpiration response to evaporative demand on chromosome 1 (position 9237408 and 8982662, respectively) mapped in intergenic regions near a gene homologous to the *SUPER APICAL DOMINANT* (*SAD1*) gene encoding an ortholog of the polymerase 1 subunit RPA34.5 that plays important roles in shoot and root development in rice (Li *et al*., 2015). Another gene coding for a heterotrimeric G protein potentially involved in the control of cell expansion via interaction with lipid metabolic pathways was identified in the region (Choudhury *et al*., 2019). Two genes encoding a TCP transcription factor and a glycosyltransferase were located at around 48 Kb upstream and downstream of the most significant SNP associated with TEr in EMMA on chromosome 4. *TCP* (*THEOSINTE BRANCHED1_CYCLOIDEA_PROLIFERATING CELL FACTOR1*) genes are involved in leaf shape in Arabidopsis (Koyama *et al*., 2010) and improved agronomic traits when overexpressed in rice (Li *et al*., 2020) while glycosyltransferase mediates the biosynthesis of prominent hemicellulose xylan (an important component of primary and secondary cell walls) in rice (Anders *et al*., 2012; Lin *et al*., 2016).

The most significant SNP on chromosome 2 (position 23902475) for transpiration response to evaporative demand is located at 9.5 Kb from a gene encoding a C2H2 zinc finger protein and 8 Kb from a gene encoding a GDSL-like lipase/acylhydrolase. The rice GDSL *BRITTLE LEAF SHEATH1* (*BS1*) gene was reported to play an important role in the maintenance of proper acetylation level on the xylan backbone (Zhang *et al*., 2017). In particular, BS1 affects secondary cell wall pattern in vessels, the *bs1* mutant having larger metaxylem pit size and reduced agronomical performances (Zhang *et al*., 2017).

## Discussion

In this study, 147 *O. glaberrima* genotypes were phenotyped in a high-throughput phenotyping platform for shoot growth and water uptake dynamics at early vegetative stage. Image-based monitoring of shoot traits (fresh weight and leaf area) and gravimetric monitoring of water loss allowed us to measure daily transpiration rate and TE at 29 DAS. Strong positive and significant correlations were observed between shoot fresh weight, total water uptake and TE in our study. Our results on *O. glaberrima* are in line with what was observed in foxtail millet (Feldman *et al*., 2018) and contradict previous claims that high water use efficiency is related to low productivity (Condon *et al*., 2002; Blum, 2009). In sorghum or pearl millet no correlation was found between TE and shoot biomass or total water use (Vadez *et al*., 2011, 2013). In fact, it appears that TE is not necessarily related to total water use or shoot growth, and the relationship between these variables depends on the species, the environments or the way water use efficiency is measured (for extensive review see Vadez *et al*., 2014). In our experimental conditions at least, it appears that larger and more vigorous *O. glaberrima* plants consume more water from the soil, but are relatively more efficient at producing biomass from that amount of water consumed. Interestingly, a significant negative correlation was observed between TE and transpiration rate, indicating that, although water loss by transpiration is higher in larger plants, transpiration per unit of leaf surface at the whole plant level is lower. We also measured the residuals of TE (TEr) that correspond in our study to the genotype-specific deviation from the relationship between biomass and water use, with genotypes deviating above the relationship being more efficient at using water than those deviating below the relationship. TEr was also significantly negatively correlated with transpiration rate, although the correlation coefficient was lower. These intriguing results suggest a stomatal regulation of transpiration rate in *O. glaberrima* genotypes with higher TE. We hypothesized that this regulation was linked to a transpiration restriction strategy in response to increasing evaporative demand.

To study the links between transpiration efficiency and transpiration response to increasing evaporative demand, we took a similar approach than Alvarez Prado *et al*. (2017) consisting in plotting daily transpiration rate with maximum reference evapotranspiration. Due to environmental conditions in the high-throughput experiment, the range of maximum evapotranspiration remained relatively low during the experiment (from 1.1 to 1.23). Still, the slope of this relationship was considered as a proxy of transpiration response to increasing evaporative demand. It was highly variable in *O. glaberrima* and under genetic control as illustrated by high broad-sense heritability. Interestingly, we observed a significant negative correlation between TE (and TEr to a lower extent) and transpiration response to increasing evaporative demand. These transpiration responses and correlations were further confirmed in a subset of genotypes where transpiration responses to much larger variation in evaporative demand (from 1.5 to 3.7) were precisely measured. Altogether, these results indicate that transpiration restriction in conditions of high evaporative demand was linked to improved TE in African rice. Transpiration response to evaporative demand is regarded as an important component trait of water use efficiency, particularly for crops growing in arid and drought-prone areas (Vadez *et al*., 2014; Shekoofa and Sinclair, 2018). In pearl millet, reducing transpiration when the evaporative demand exceeds a certain threshold allowed water conservation during the vegetative growth that could be used at reproductive stage for better yield (Vadez *et al*., 2013). Early vigor accompanied by increased TE through transpiration restriction strategies may be particularly advantageous for upland rice genotypes growing in rainfed agroecosystems, especially when competing against weeds or under the occurrence of a drought stress.

Exhaustive measurements of transpiration profile under changing temperature and relative humidity over the course of the day allowing precise measurement of the transpiration restriction phenotype has often been low throughput (Gholipoor *et al*., 2010, 2013; Jyostna Devi *et al*., 2010; Jauregui *et al*., 2018). To our knowledge, our study is pioneer in reporting measurements of transpiration restriction in a crop at a throughput compatible with association genetic analyses. Recent development of an imaging platform combined with lysimetric capacity allowing monitoring of transpiration response to high VPD in natural conditions (Vadez *et al*., 2015) and automation of transpiration profile features generation (Kar *et al*., 2020) will be instrumental for high-throughput phenotyping of plant water use-related traits and identification of their genetic determinants with breeding perspectives.

Roots are the primary sites of water uptake and play important roles in maintaining whole plant water status, balancing water acquisition and water flow to match shoot water demand (Maurel *et al*., 2010; Vadez, 2014). Root traits controlling radial root conductance including aquaporin functions (Reddy *et al*., 2017; Sivasakthi *et al*., 2017, 2020; Grondin *et al*., 2020) or apoplastic barriers (Calvo-Polanco *et al*., 2021; Reyt *et al*., 2021) as well as those controlling root axial conductance including metaxylem diameter (Richards and Passioura, 1989) have been linked to water balance and plant transpiration efficiency in several crops. In this study, we observed a positive significant correlation between root to shoot ratio and transpiration response to increasing evaporative demand. Assuming that root dry weight is largely related to root surface, these results indicate that the balance between root and shoot surfaces are important for transpiration response to increasing evaporative demand. Plants with larger root surface as compared to shoot surface appeared less sensitive to the increase in evaporative demand, possibly due to the ability of the root system to maintain water acquisition in response to the increased water demands by the shoots. Our results therefore suggest that decreased carbon allocation towards the roots, and possibly decreased root surface may be beneficial for the transpiration restriction phenotype. Further investigations are needed to determine the contribution of root architectural, anatomical or physiological traits in the transpiration restriction phenotype and how these relate to drought tolerance in *O. glaberrima*.

Interestingly, our GWAS approach confirmed the importance of roots in the control of TE and transpiration response to increasing evaporative demand. Indeed, we identified several genetic regions associated with these traits that contain genes potentially involved in root development or water transport. In particular, a cluster of aquaporin genes was located near the association for TE observed on chromosome 7, amongst which LOC_Os07g26660 appears root-specific. These genes encode type-2 Plasma membrane Intrinsic Proteins that are known to play important roles in root radial water transport. Their expressions have also been associated with the control of TE and transpiration restriction as they may quickly regulate root water flow to respond, or not, to changes in transpiration when the evaporative demand is increasing (Shekoofa and Sinclair, 2018). Aquaporins function also have important roles in root and shoot growth coordination (Ehlert *et al*., 2009; Maurel *et al*., 2010). In fact, this GWAS association on chromosome 7 was also found for shoot fresh weight, although just below the significance threshold.

A strong marker-trait association for TE and shoot fresh weight at 29 DAS was located on chromosome 5. This association collocated with a known QTL for early vigor identified in *O. sativa* (Cui *et al*., 2002). This suggests that this QTL for early vigor is conserved in Asian and African rice. An interesting candidate gene coding for a polyprenyl synthetase protein potentially involved in plastoquinone biosynthesis and more generally in photosynthesis was located close to the most significant SNP. This suggest that a more efficient photosynthetic machinery might be responsible for increased early vigor. Further work will be needed to test this exciting hypothesis.

TEr and transpiration response to increasing evaporative demand shared an association on chromosome 1 where a candidate gene involved in cell wall biosynthesis was identified. This result confirms the links between these two traits and support the hypothesis that transpiration restriction is an important component of TE in *O. glaberrima*. Another candidate gene encoding a GDSL protein possibly involved in cell wall biosynthesis was identified in close vicinity of the most significant association on chromosome 2 for transpiration response to increasing evaporative demand (Zhang *et al*., 2017). Cell wall properties potentially play important roles in the apoplastic water path in roots and shoots. This path, complementary to the symplasmic path (from cell to cell through aquaporins or plasmodesmata), is supposedly predominant under increasing evaporative demand, i.e. under conditions where transpiration restriction occurs (Tharanya *et al*., 2018; Sivasakthi *et al*., 2020). The effects of cell wall content and mechanical properties on plant water transport properties have been poorly studied. Simulations using a model coupling water fluxes and cell wall mechanics recently suggested that heterogeneities in cell wall mechanical parameters in tissues impacted water flow and growth rate (Cheddadi *et al*., 2019). Moreover, affecting cell wall composition had significant effects on xylem vessel wall patterning in rice, which may further impact axial water flow (Zhang *et al*., 2017).

In conclusion, high-throughput phenotyping of water use-related traits in *O. glaberrima* showed that transpiration restriction under increasing evaporative demand may be an important strategy to improve TE in *O. glaberrima* rice, which is partly controlled by the balance between root and shoot growth. The functional mechanisms of such control in terms of water fluxes are still unknown although association genetics pointed to mechanisms linked to cell wall composition and patterning.

## Supplementary data

**Supplementary Table S1:** Analysis of variance for shoot fresh weight (SFW) measured at 29 days after sowing in *O. glaberrima* in the large-scale experiment.

**Supplementary Table S2:** Analysis of variance for total water uptake (TWU) measured at 29 days after sowing in *O. glaberrima* in the large-scale experiment.

**Supplementary Table S3:** Analysis of variance for transpiration efficiency (TE) measured at 29 days after sowing in *O. glaberrima* in the large-scale experiment.

**Supplementary Table S4:** Analysis of variance for residuals of transpiration efficiency (TEr) measured at 29 days after sowing in *O. glaberrima* in the large-scale experiment.

**Supplementary Table S5:** Analysis of variance for transpiration rate (TR) measured at 29 days after sowing in *O. glaberrima* in the large-scale experiment.

**Supplementary Table S6:** Analysis of variance for transpiration response to increasing evaporative demand (SlopeTR) measured in *O. glaberrima* in the large-scale experiment.

**Supplementary Table S7:** Transpiration response (SlopeTR) and inflexion in transpiration rate (InflexionTR) under increasing evaporative demand measured in the subset of *O. glaberrima* genotypes in the small-scale experiment.

**Supplementary Fig. S1**. Regression model used to estimate shoot fresh biomass and leaf area based on ground truth measurements.

**Supplementary Fig. S2:** Residuals of transpiration efficiency.

**Supplementary Fig. S3**. Histograms, QQ-plots, and plots of residuals against fitted or index values for fixed (fix) or random (ran) genotypic effects on shoot fresh weight measured at 29 days after sowing in the large-scale experiment.

**Supplementary Fig. S4:** Histograms, QQ-plots, and plots of residuals against fitted or index values for fixed (fix) or random (ran) genotypic effects on total water uptake measured at 29 days after sowing in the large-scale experiment.

**Supplementary Fig. S5:** Histograms, QQ-plots, and plots of residuals against fitted or index values for fixed (fix) or random (ran) genotypic effects on transpiration efficiency measured at 29 days after sowing in the large-scale experiment.

**Supplementary Fig. S6:** Histograms, QQ-plots, and plots of residuals against fitted or index values for fixed (fix) or random (ran) genotypic effects on residuals of transpiration efficiency measured at 29 days after sowing in the large-scale experiment.

**Supplementary Fig. S7:** Histograms, QQ-plots, and plots of residuals against fitted or index values for fixed (fix) or random (ran) genotypic effects on transpiration rate measured at 29 days after sowing in the large-scale experiment.

**Supplementary Fig. S8:** Water use-related traits in the subset of *O. glaberrima* genotypes.

**Supplementary Fig. S9:** Correlation between water use-related traits measured in the large-scale experiment at 29 days after sowing (PhenoArch) and in the small-scale experiment at 35 days after sowing (Subset).

**Supplementary Fig. S10:** Transpiration response to increasing evaporative demand.

**Supplementary Fig. S11:** Histograms, QQ-plots, and plots of residuals against fitted or index values for fixed (fix) or random (ran) genotypic effects on transpiration response to increasing evaporative demand (SlopeTR) measured in the large-scale experiment.

**Supplementary Fig. S12:** Transpiration response to increasing evaporative demand in the subset of *O. glaberrima* genotypes.

**Supplementary Fig. S13:** Correlation between water use-related traits and plant morphology in a subset of *O. glaberrima* genotypes.

**Supplementary Fig. S14:** Genome wide association studies for shoot fresh weight, total water uptake, transpiration efficiency (TE), and residuals of transpiration efficiency (TEr) in *O. glaberrima*.

**Supplementary Fig. S15:** Genome wide association studies for transpiration response to increasing evaporative demand in *O. glaberrima*.

**Supplementary Fig. S16:** Repartition of shoot fresh weight according to the allelic version at the most significant SNP (Chr5_26971730) for the GWAS association on chromosome 5.

## Acknowledgments

This work was supported by the Institut de Recherche pour le Développement, the CGIAR Research Program (CRP) on rice-agrifood systems (RICE, 2017-2022) and the Agence Nationale de la Recherche (grants ANR-17-MPGA-0011 to VV). Financial support by the Access to Research Infrastructures activity in the Horizon 2020 Programme of the EU (EPPN^2020^ Grant Agreement 731013) is gratefully acknowledged. PA was supported by a doctoral fellowship from the French Ministry of Higher Education. BEE was supported by the Centre National de la Recherche Scientifique et Technologique of Gabon. The authors acknowledge the IRD iTrop HPC (South Green Platform) at IRD Montpellier for providing HPC resources (https://bioinfo.ird.fr – http://www.southgreen.fr). We are also grateful to Harold Chrestin and Laurence Albar (IRD) for providing seeds of the *O. glaberrima* genotypes, to Gabriel Quiroga García (CISC, Spain), Romane Le Roy (INRAE, France) and all the PhenoArch staff for the technical support, to Emilie Millet (Wageningen University, Netherlands) for her kind support in the PhenoArch data analyses, to all the *Cereal Root Systems* team of UMR DIADE and to Soumyashree Kar (Indian Institute of Technology, India) for their kind support on the data analysis of the Phenospex experiment.

## Author contributions

LCB, CW, AC, PG, AGD, BM, RA, VV, LL, PC and AG designed the study. PA, BEE, DM, MSN, MP, NL and LCB performed the experiments with help from all co-authors. PA, BEE, MSN, NL, RP, LCB, VV, LL, PC and AG analysed the data. AG, PC, LL and VV wrote the first draft of the manuscript that was edited and approved by all co-authors.

## Data availability

The data supporting the findings of this study are available from the corresponding author upon request.

